# A comprehensive analysis of the inherited lncRNA and circRNA repertoire of zebrafish

**DOI:** 10.1101/2024.12.31.630970

**Authors:** Dheeraj Chandra Joshi, Aakanksha Kadam, Chetana Sachidanandan, Beena Pillai

**Affiliations:** CSIR-Institute of Genomics and Integrative Biology, New Delhi, 110025, India; Academy of Scientific and Innovative Research (AcSIR), Ghaziabad 201002, India

**Keywords:** Epigenetic inheritance, lncRNA, circRNA, zebrafish

## Abstract

Inherited non-coding RNAs can be the third major component of epigenetic information transfer from one generation to the next. Here, we present a comprehensive resource of lncRNAs and circular RNAs that are inherited, compiled from meta-analysis of zebrafish transcriptomics data and comparative genomics with mouse and human. Maternal and paternal inheritance of mRNA into the zygote is accepted to be an important regulator of embryonic development as well as adult characteristics. Although inheritance of certain specific miRNAs is known, other non-coding RNA inheritance remains less explored. We performed a comprehensive analysis of the inherited lncRNAs and circRNAs in zebrafish. We discovered that nearly 20% of all known lncRNA and 7% of circRNAs are inherited. Many of these lncRNAs are conserved in mammals, and are expressed widely in adult tissues of zebrafish. The male and female gametes carry a highly similar pool of inherited lncRNAs, with only a few sperm/ oocyte specific transcripts. The majority of inherited circRNAs originate from genes important for fertilization and can potentially regulate translational processes. Contrary to general belief, the inherited lncRNAs and circRNAs do not undergo degradation en masse coincidental to zygotic genomic activation, suggesting that these RNAs may have more sustained roles in development.

## Introduction

The genetic information encoded by DNA can be regulated and transmitted through non-DNA-based means collectively termed epigenetic inheritance. Such mechanisms play crucial roles in cellular function, embryonic development and adaptation. Besides the parental DNA, the zygote is also known to inherit specific DNA methylation patterns, histone modification patterns, proteins, mRNAs and regulatory RNAs that influence how the genome is read and interpreted. These factors can have housekeeping or developmental roles and could be involved in the adaptation to changing environmental conditions and stressors. Small RNA inheritance has been particularly well studied in the *C. elegans*. Inherited endo-siRNAs, miRNAs and piRNAs in *C. elegans* play important roles in processes as diverse as maintaining genome integrity(Lee et al. 2012), clearance of maternal transcripts(Rouget et al. 2010), adaptation to pathogens(Kaletsky et al. 2020), and starvation(Rechavi et al. 2014) We have previously shown the inheritance of miR-34 in zebrafish(Soni et al. 2013) and subsequently others have shown that reduced levels of miR-34/449 in the mouse sperm is crucial for the inheritance of chronic social instability stress associated phenotype(Champroux et al. 2024). Parental inheritance of other types of small RNA has also been shown, for instance tRFs (tRNA fragments) or tsRNAs (tRNA derived small RNAs) comprising 60-70%, dominate the small RNA repertoire of mouse sperm(Conine et al. 2018). Additionally, mouse sperm also contain miRNAs (∼10-20%) and they have been implicated in both normal embryonic development as well as inheritance of traits. microRNAs such as miR-880, miR-17-92 and miR-106b-25 clusters, and miR-34b/c inherited from mouse sperm are shown to be important for embryo viability(Conine 2019).

Long regulatory RNAs such as lncRNAs and circRNAs are emerging as important components of epigenetic regulation. High throughput sequencing datasets have revealed the presence of hundreds to thousands of lncRNAs in gametes and zygote of diverse organisms including mouse(Zhang et al. 2017; Karlic et al. 2017), pig(Yang et al. 2022) and boar(Fraser et al. 2020), cattle(P. Wang, Paquet, and Robert 2023; J. Wang, Koganti, and Yao 2020), ram (Hitit, Kaya, and Memili 2024) and humans(Zhang et al. 2019; Corral-Vazquez et al. 2021). Some studies have also reported an association between sperm lncRNA profile and fertility in ram (Hitit, Kaya, and Memili 2024) and humans(Zhang et al. 2019). One recent study in mice showed altered lncRNA profile in sperm in response to stress hormone and injecting the altered lncRNAs in normal zygotes resulted in behavioral phenotypes in the offspring (Hoffmann et al. 2024).

Circular RNAs (circRNAs) are covalently closed RNA molecules that are resistant to exonuclease action. Since their discovery around 40 years ago, numerous circRNAs have been identified in gametes and zygote of various organisms including pig(Cao et al. 2019) (Cao, Gao et al. 2019), humans and mouse(Ragusa et al. 2019)(Ragusa, Barbagallo et al. 2019). Some studies have also reported the presence of hundreds of circRNAs in pre-implantation embryos of humans (Dang et al. 2016)(Dang, Yan et al. 2016) and mouse (Fan et al. 2015)(Fan, Zhang et al. 2015). circRNAs are associated with male infertility (Manfrevola et al. 2020)(Manfrevola, Chioccarelli et al. 2020,(Tang et al. 2023)) and ovarian maturation(J. Li et al. 2021).

The presence of lncRNAs and circRNAs in gametes of all these species suggests that their inheritance in the zygote could be a widespread phenomenon. Unlike the inherited small non-coding RNAs, the inheritance of lncRNAs and circRNAs has not been well studied. The ex-utero fertilization of zebrafish makes them an excellent model to study RNA inheritance. Extensive knowledge of zygotic genome activation and early development in zebrafish lends itself to functional studies of inherited RNAs.

Here, we have identified the inherited lncRNA and circRNA repertoire of zebrafish and explored their features. We identified 2093 lncRNA and 270 circRNA that are inherited in zebrafish. The genes neighboring to the inherited lncRNAs are enriched for translation and embryonic development related pathways. Majority of inherited lncRNAs and circRNAs are present in both gametes. There is a distinct population of inherited lncRNA and circRNAs that are exclusively of maternal origin. We find that more than 250 inherited lncRNAs of zebrafish have at least one conserved counterpart in the inherited lncRNA pool of both mouse and human. Notably, we found that the majority of inherited lncRNAs and circRNAs retain their zygotic expression till 4 hours post-fertilization (hpf) and there is no ZGA-associated degradation countering the idea that there is a significant turnover and replacement of RNA population around the ZGA phase.

## Results

### 20% zebrafish lncRNAs are inherited

To identify inherited lncRNAs and circRNAs in zebrafish, we utilized the publicly available RNA sequencing datasets of gametes (sperm and oocyte) and zygote (Supplementary file 1). The datasets were reanalyzed at transcript level using the latest zebrafish assembly, GRCz11 (Figure 1A). We reasoned that a transcript detectable in zygote is likely to be inherited since the zygote is known to be transcriptionally inactive. To bolster our conclusion, we also checked whether these zygotic transcripts are detected in at least one of the gametes. The transcripts passing these criteria were considered to be inherited.

**Figure 1.**
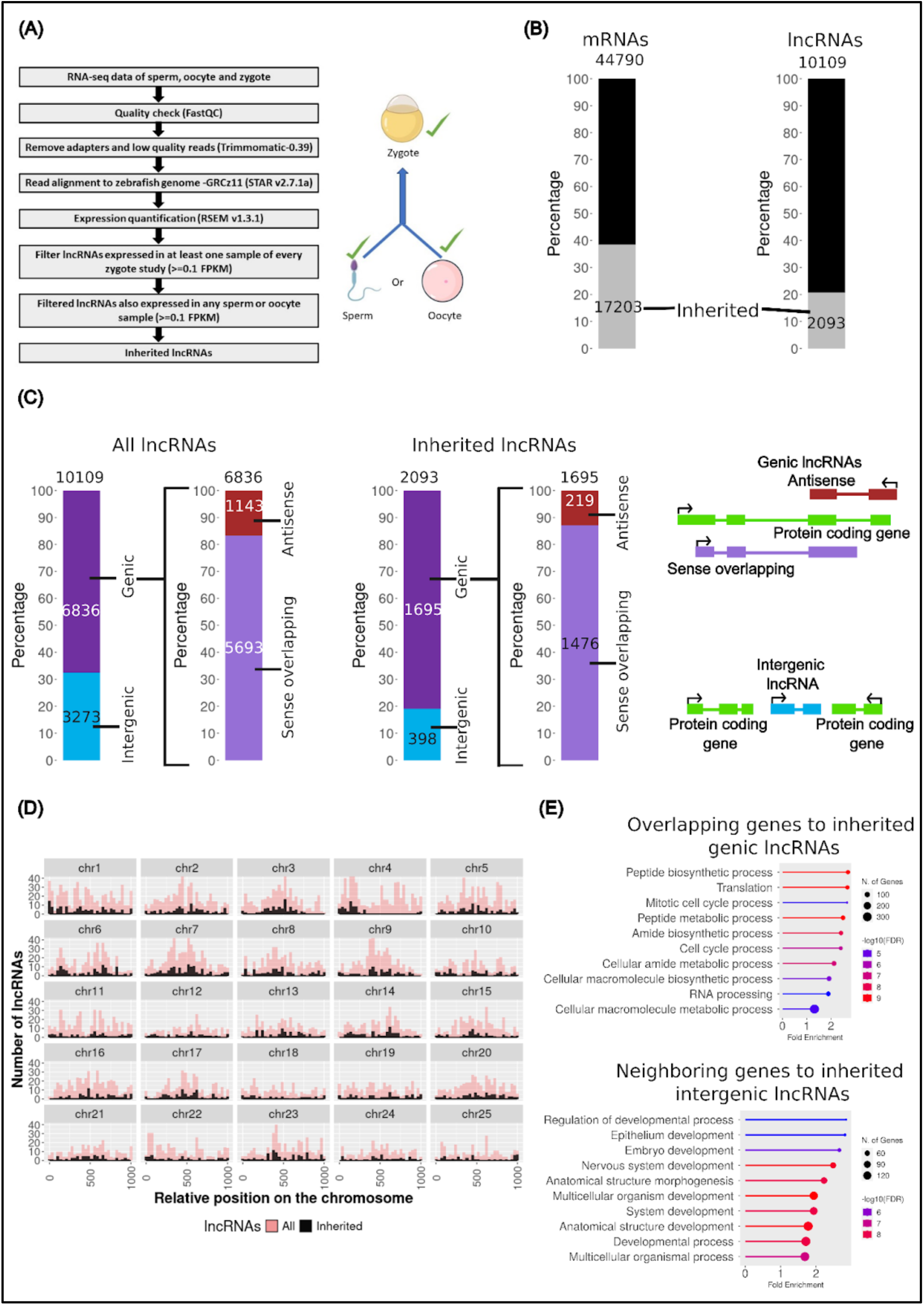
Around 20% of lncRNAs in zebrafish are inherited. Inherited lncRNAs arise from all chromosomes and are mostly located in the vicinity of metabolic and developmental genes. (A) Strategy used to identify the inherited lncRNAs in zebrafish (B) Stacked bar plots showing the percentage of inherited lncRNAs and mRNAs (C) Stacked bar plots showing the subcategories of lncRNAs. Above the stacked bar plot the total number of lncRNAs are written and the numbers for subcategories are written on the top of individual categories. The schematics of the different types of lncRNAs are shown on the right of the stacked bar plots. (D) Density plot showing the genomic loci of inherited lncRNAs. The 25 chromosomes of zebrafish are scaled to the same length with zero representing the start of the chromosome and 1000 representing the end of the chromosome. Overall density of all known lncRNAs (10109) is shown in the background (Pink) and the density of inherited lncRNAs (2093) is shown in the foreground (Black). (E) GO analysis of protein coding genes overlapping or flanking to genic and intergenic inherited lncRNAs.

We found that approximately 20% of all known lncRNAs in zebrafish are inherited, which amounts to 2093 inherited lncRNAs out of 10109 total lncRNAs (Figure 1B). In contrast, approximately 40% of all mRNAs are inherited. Out of the 2093 inherited lncRNAs identified in this study, the majority (∼80%) are genic lncRNAs, whereas the rest are intergenic (Figure 1C; Supplementary file 2). In this study, we use the term ‘genic’ to be synonymous to protein-coding loci. The percentage of inherited genic lncRNAs (80%) is greater than the total percentage of genic lncRNAs (65%) showing a bias towards coding regions in the inherited pool. The genic lncRNAs are further subcategorized into antisense and sense overlapping lncRNAs.

Genomic locations of lncRNAs and the neighboring protein coding genes can reveal their likely function as several lncRNAs are known to have cis-regulatory effects on the genomic loci of their origin. To explore whether the inherited lncRNAs arise from select locations, like telomeres or heterochromatin in the genome, we visualized the density of inherited lncRNAs versus the overall density of lncRNAs on each of the zebrafish chromosomes. We found that inherited lncRNAs arise almost uniformly from all over the genome (Figure 1D) with the exception of two notable loci in chr4 and chr25. These genomic regions spanning several megabases do not give rise to any inherited RNAs, even as they contain many lncRNA and protein-coding genes. To understand what category of genes are enriched in the protein coding genes from the inherited lncRNAs loci, we performed a Gene Ontology (GO) analysis at “Biological Processes” level for genic and intergenic lncRNAs. We found that protein coding genes from the vicinity of genic inherited lncRNA genes are mainly involved in processes like metabolism and cell cycle. On the other hand, protein coding neighbors of the intergenic inherited lncRNAs loci are majorly enriched for specific developmental processes (Figure 1E). Both categories of genes are crucial for early development and inherited lncRNAs may be the regulators of these genes.

### Inherited lncRNAs correspond to broadly expressed zygotic lncRNA

One of the typical features associated with the lncRNAs is that they are low expression transcripts as compared to protein coding transcripts (mRNAs). We plotted the zygotic expression of all the inherited lncRNAs and inherited mRNAs in the form of violin plots. Consistent with the generalization regarding low expression of lncRNAs, we found that inherited lncRNAs have a four-fold lower median expression than inherited mRNAs in the zygote (Figure 2A). However, the expression range of inherited lncRNAs is very broad and a minority of inherited lncRNAs (359/2093) are present at levels higher than the median mRNA expression in the zygote. Also, we found that there is some variation in the expression levels among the inherited lncRNA subcategories (antisense, sense overlapping and intergenic). The median expression for sense overlapping inherited lncRNAs is the highest and approximately 2-fold higher than the antisense and intergenic inherited lncRNAs (Figure 2B). We have also identified the presence of some notable inherited lncRNAs (Figure 2B) e.g. several inherited antisense lncRNAs are derived from hox loci and many sense overlapping inherited lncRNAs are derived from ribosomal protein genes (rpl/rps genes) (Supplementary file 3). In the intergenic category we found two well-known lncRNAs; MALAT1, traditionally studied for its role in cancer and Cyrano, previously shown experimentally to be inherited in zebrafish by our group (Sarangdhar et al. 2018).

**Figure 2.**
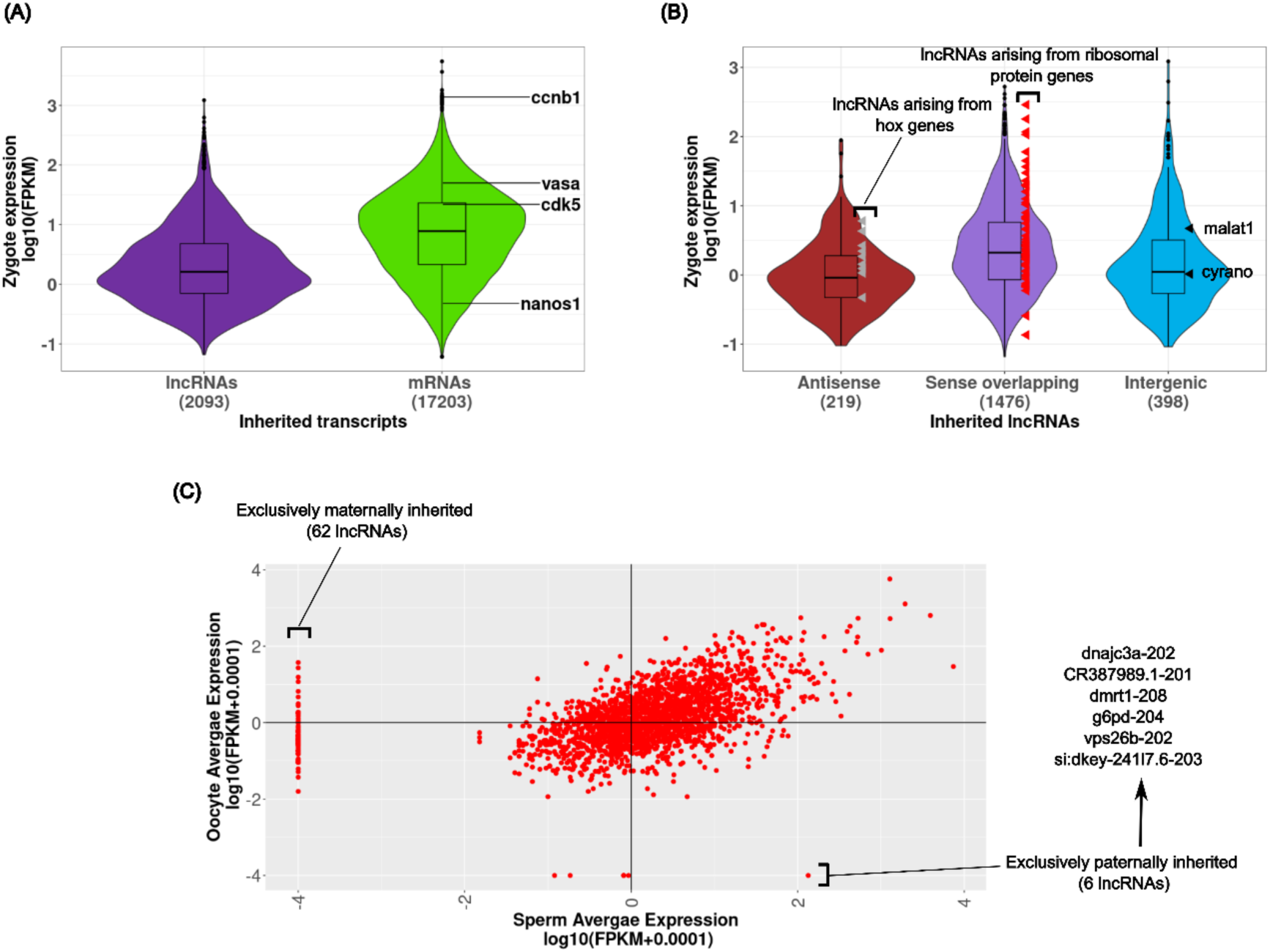
Inherited lncRNAs have a broad expression range in the zygote. Inherited lncRNAs show similar expression and diversity in sperm and oocytes. (A) Violin plots showing the expression of all inherited lncRNAs (2093) and inherited mRNAs (17203) in the zygote. The expression levels of well-known inherited mRNAs have been marked. (B) Violin plots showing the expression of antisense (219) sense overlapping (1476) and intergenic inherited mRNAs (398) in the zygote. The expression levels of some notable inherited lncRNAs in each category are marked. (C) Scatter plot showing the expression levels of 2093 inherited lncRNAs in the sperm and oocytes.

### Sperm and oocytes contain similar lncRNA pool

Traditionally, sperm was considered to be merely a vehicle for paternal DNA during fertilization and it was believed that all the non-DNA factors were maternally supplied. However, recent studies have shown that sperm-derived miRNAs and tsRNAs were delivered to the zygote (Chen et al. 2016)(Q. Chen et al. 2016). To understand whether the inherited lncRNAs in zebrafish are sperm-derived or oocyte-derived, we plotted the expression of 2093 inherited lncRNAs in sperm and oocyte in the form of a scatter plot. We found that the majority of inherited lncRNAs are expressed in both sperm and oocyte at similar levels (Figure 2C). We also found 62 oocyte specific and 6 sperm specific inherited lncRNAs (Figure 2C; Supplementary file 4). The six exclusively paternally inherited lncRNAs overlap with protein coding loci, including one arising from the Dmrt1 gene locus, which is previously known to be involved in transcriptional regulation during germ cell lineage commitment. The most abundant oocyte specific lncRNA, overlaps with the gene Csde1, previously implicated in brain development.

### Post-fertilization stability of inherited lncRNAs is maintained until ZGA, with broad expression observed across multiple adult tissues

Next, we wanted to understand the functional relevance of inherited lncRNAs. One of the possibilities is that inherited lncRNAs could simply be carried over following gametogenesis with no specific function in the embryo. Alternatively, they could act as a source of nucleotides in the early embryo following their degradation. Yet another and more exciting possibility is that the inherited lncRNAs might be regulators of specific zygotic genes. To explore whether the inherited lncRNAs are retained post-fertilization or degraded, we analyzed the publicly available RNA sequencing datasets that included the post fertilization stages of zebrafish development (0 hpf, 1 hpf, 2 hpf, 3 hpf and 4 hpf). The expression patterns of 2093 inherited lncRNAs could be separated into 4 distinct groups (Figure 3A; Supplementary file 5). It was observed that the majority (∼96%) of inherited lncRNAs lie in group 1-3 that mostly retain their original expression till 4 hpf and only ∼4% of lncRNAs in group 4 show patterns consistent with degradation. This observation suggests that the majority of lncRNAs are not degraded, at least until ZGA and thus unlikely to be used as a nucleotide source.

**Figure 3.**
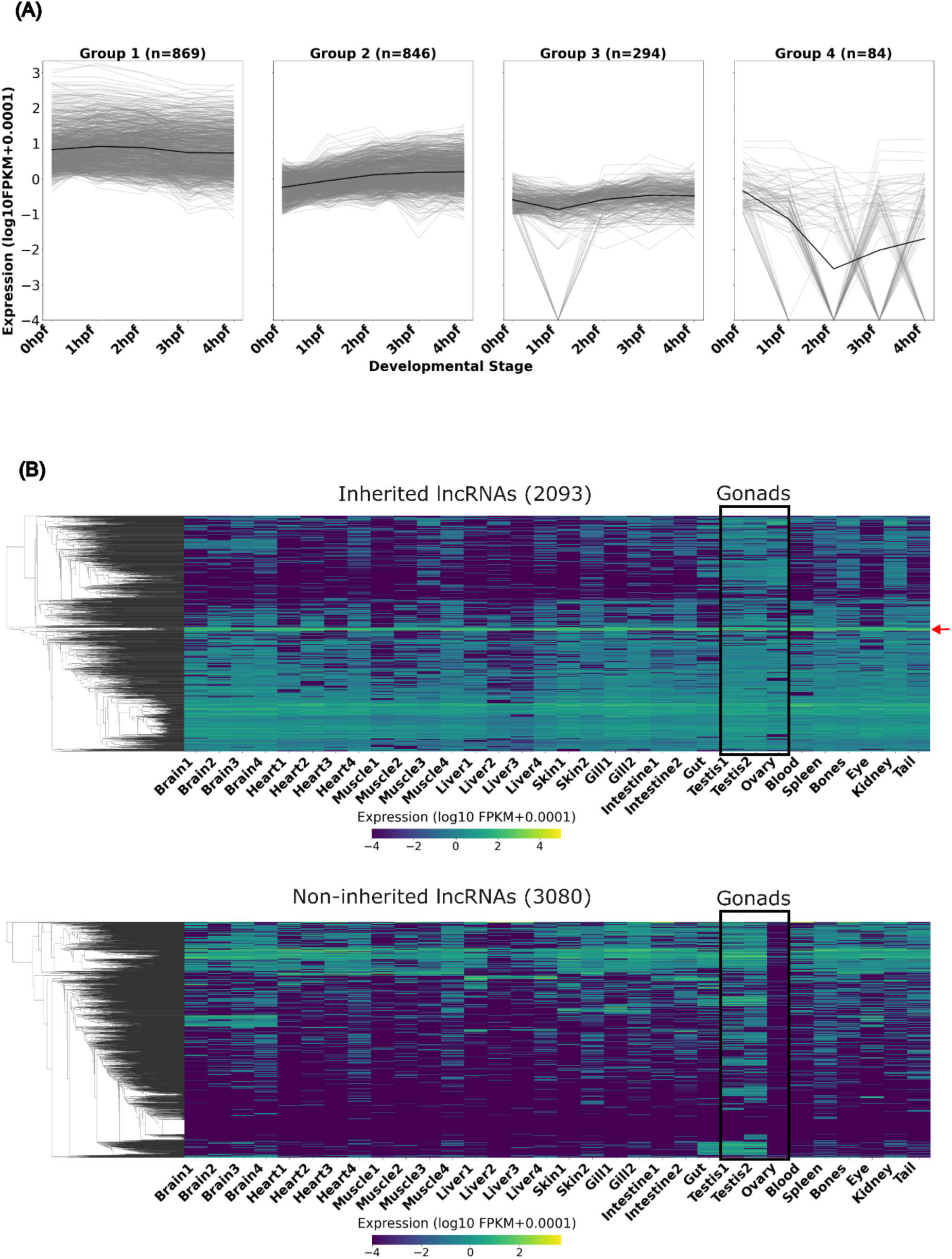
Inherited lncRNAs are mostly stable post fertilization till ZGA. Inherited lncRNAs are broadly expressed across multiple adult tissues. (A) Line graphs showing the expression dynamics of inherited lncRNAs post fertilization. Individual lncRNA expression profiles are shown in light gray lines and the average expression trend for each group is highlighted in a thick black line. The groups are arranged by the number of lncRNAs assigned to each group (group 1 -highest number to group 4 -lowest number). hpf-hours post fertilization. (B) Clustered heatmap showing the expression levels of inherited lncRNAs (2093) and non-inherited lncRNAs (3080) in adult tissues of zebrafish. The red arrow highlights a set of inherited lncRNAs derived from ribosomal protein genes that are consistently expressed in all the tissues.

Also, we explored the expression pattern of inherited lncRNA counterparts in adult tissues of zebrafish to understand their specificity and possible functional relevance later in life. We utilized the bulk RNA sequencing datasets of 14 different tissues from 4 different research groups for this analysis (Supplementary file 1). As expected, we found the expression of inherited lncRNAs to be enriched in the gonads (testis and ovary) (Figure 3B). Unexpectedly, we also found that the majority of inherited lncRNAs show a broad tissue expression pattern that was sharply in contrast with the non-inherited lncRNAs that show much more tissue specificity (Figure 3B; Supplementary file 6).

### Over 250 zebrafish inherited lncRNAs have conserved counterparts in both mouse and human zygotes

To find if the zebrafish inherited lncRNAs have conserved counterparts in mouse and human zygotes, we first identified the inherited lncRNAs in mouse and human using a similar criteria as we used for zebrafish. We found 3785 (∼7%) inherited lncRNAs in the mouse and 4120 (∼4%) inherited lncRNAs in humans (Supplementary file 7). The details of the publicly available RNA sequencing datasets used for this analysis are provided in the methods section (Supplementary file 1). One of the general features associated with lncRNAs is poor sequence conservation. However, syntenic conservation, by virtue of vicinity or overlap to conserved protein coding genes, is relatively more common. To identify the inherited lncRNAs in zebrafish that may be conserved with mouse and human, three different definitions for conservation were used; sequence conservation, conservation of overlap with protein coding genes and syntenic loci conservation (Figure 4A). We found that 276/ 2093 inherited lncRNAs of zebrafish have a conserved and inherited counterpart in both mouse and human (Supplementary file 8). Consistent with the generalization, we found only 26 out of 2093 zebrafish inherited lncRNAs to be conserved by sequence (Figure 4B). The sequence identity ranged from 75% to 95% and query coverage ranged from 43 to 745 nucleotides. Majority of conserved inherited lncRNAs (250/276) were exclusively conserved by synteny (overlap conserved or loci conserved). One lncRNA can be conserved by more than one definition and the Venn diagram shows such overlap between the three conservation categories (Figure 4B).

**Figure 4.**
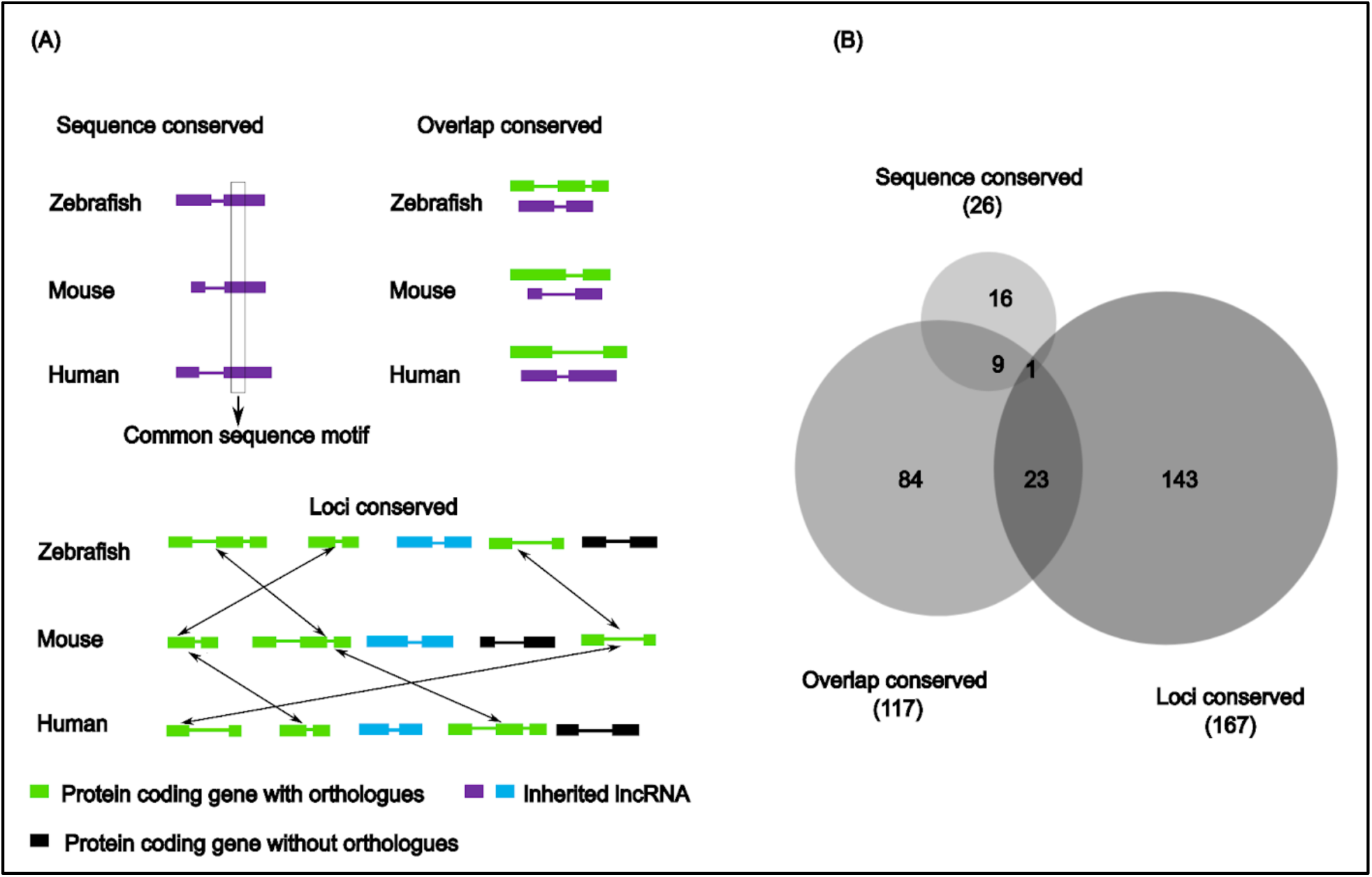
Counterparts of more than 250 zebrafish inherited lncRNAs are present in both mouse and human zygotes. (A) Schematic showing different strategies used to identify the conserved counterparts of zebrafish inherited lncRNAs in mouse and human. (B) Venn diagram showing the overlap between the three conservation categories.

### 7% of the total identified circRNAs, spread across the genome, are inherited in zebrafish

To explore zebrafish circular RNAs in gametes and zygote, we reanalyzed publicly available RNA sequencing datasets of oocytes, sperm and zygote from NCBI GEO/SRA database (Supplementary file 1). CircRNA detection tool CIRI2 (Gao, Zhang, and Zhao 2018) (Gao, Zhang, and Zhao 2018) was used to identify 3678 circRNAs. A circRNA was called only if at least 2 back splice reads overlapping with the junction were supporting the junction and with minimum of 0.05 junction ratio, which is indicative of the expression of a circRNA relative to its linear RNA counterpart (Figure 5A). A candidate circular RNA was considered inherited only if it was expressed in at least one sample of the zygote, as well as in either or both of the gametes. We identified a total of 270 inherited circRNAs in zebrafish (Figure 5B; Supplementary file 9). Among them ∼86% (231) of these were of genic origin, aligned to an annotated protein coding gene and ∼14% (198) came from intergenic regions, likely non-coding RNA loci (Figure 5B). A circRNA was termed as genic, if even one of its coordinates were located in any annotated gene, irrespective of the location of the other end. Multiple back splice junctions were identified from a single host gene.

**Figure 5.**
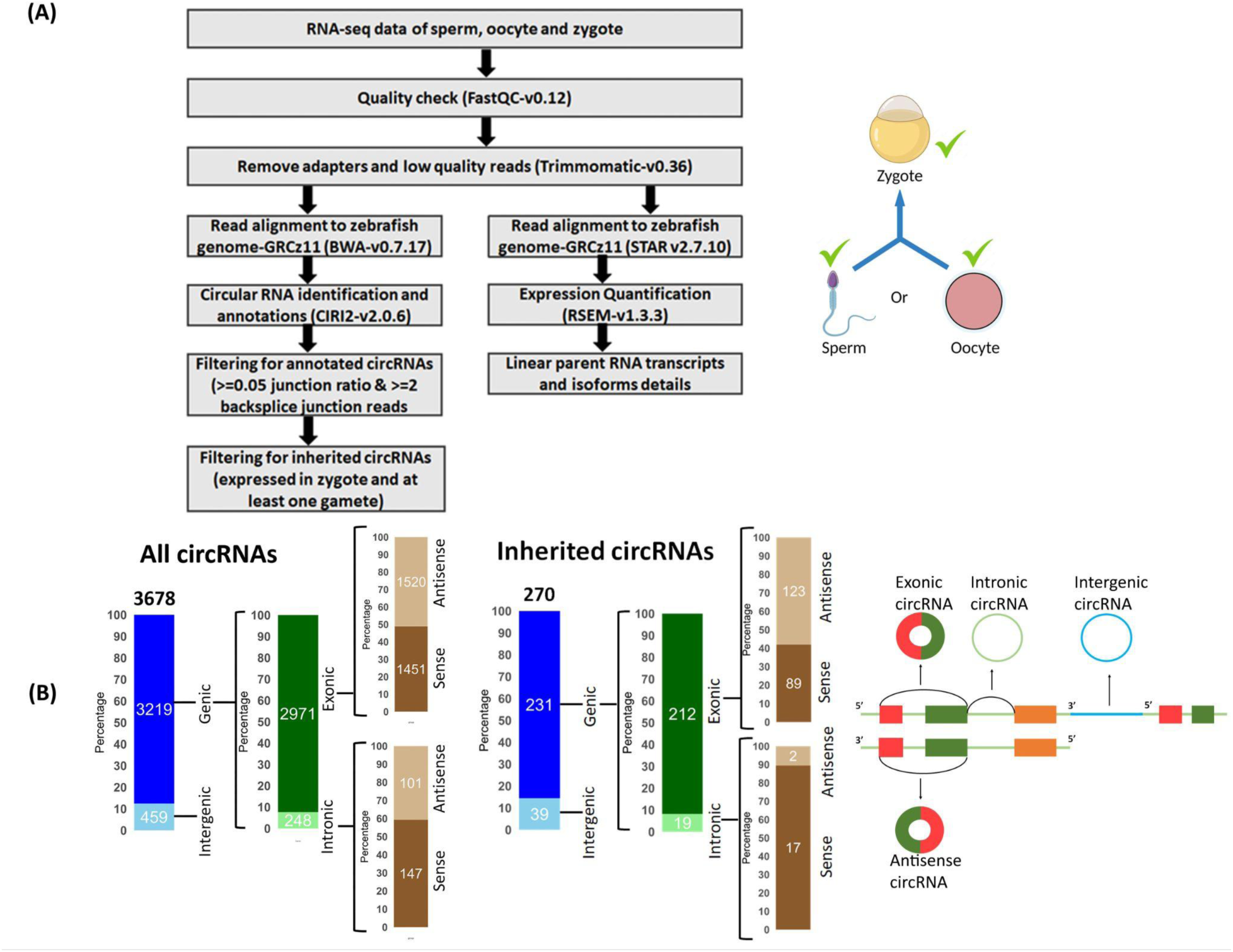
Around 7% of circRNAs in zebrafish are inherited. (A) Schematic description of bioinformatics analysis for discovery of inherited circular RNAs in zebrafish. (B) Stacked bar plots representing the amount of exonic, intronic and intergenic circular RNAs identified in all circRNAs and inherited circular RNAs. Above the stacked bar plot the total number of circRNAs are written and the numbers for subcategories are written on the top of individual categories. The schematics of the different types of circRNAs are shown on the right of the stacked bar plots.

### Inherited CircRNAs originate from genes involved in fertilization

We mapped the density of inherited circRNAs origins across the genome and found that chromosome 3 had the highest number of circRNA loci per million bases across its length (Figure 6A). circRNAs often regulate the gene of their origin (Liang et al. 2019)(Liang, Wong et al. 2019, (Liu et al. 2021). To identify the category of genes that generate inherited circRNAs we performed a GO analysis at ‘biological processes’ level for the genic inherited circRNAs. We found that the majority of the host genes were involved in the process of fertilization such as acrosome reaction, egg coat formation and zona pellucida interaction with sperm (Figure 6B).

**Figure 6.**
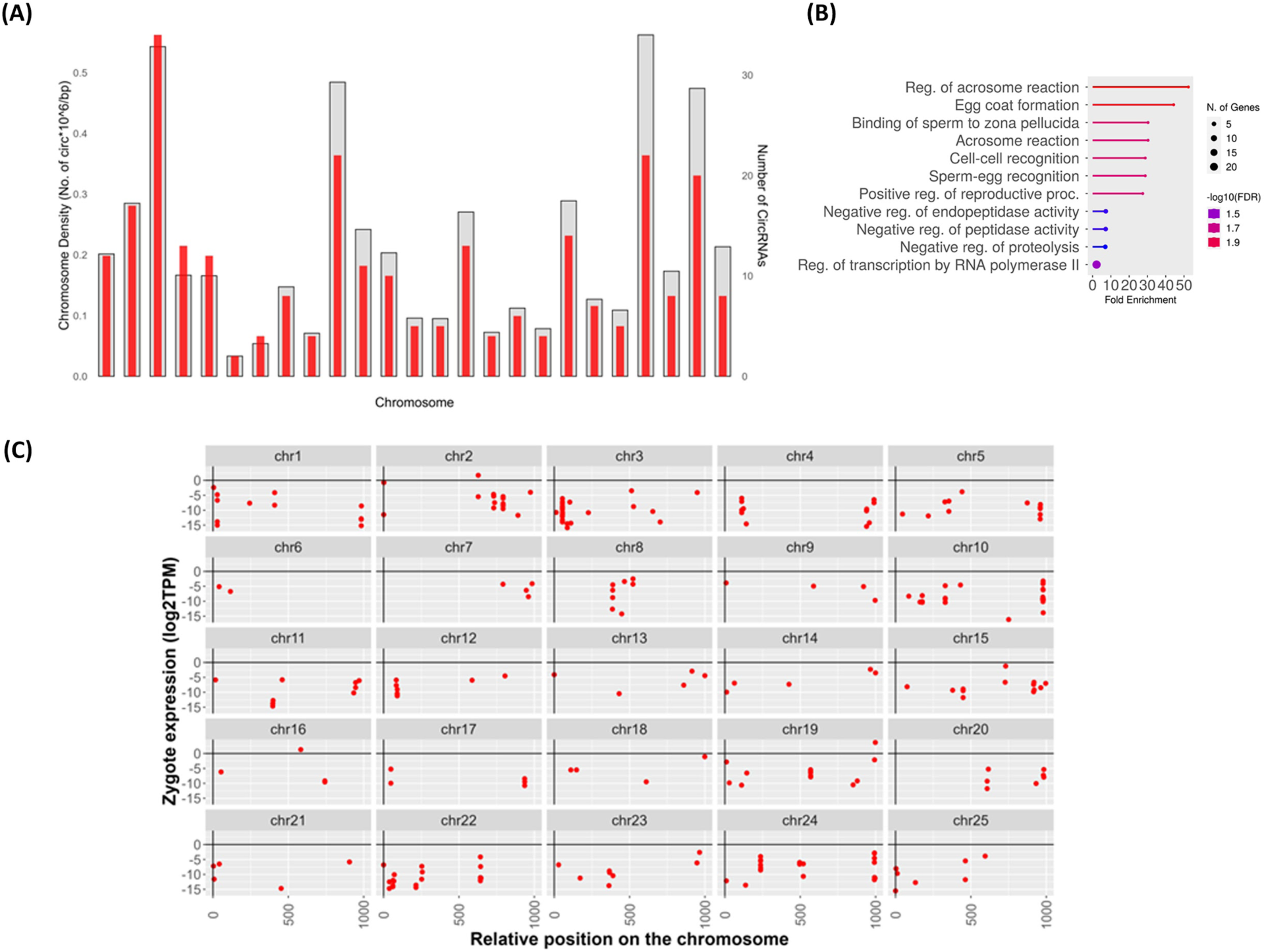
Inherited circRNAs originate from all chromosomes. (A) Density-wise distribution of inherited circRNAs across chromosome length. Overall density of inherited circRNAs is shown in the background (Grey) and the number of inherited circRNAs is shown in the foreground (Red). Chromosomes are arranged in order 1 to 25 from left to right. (B) GO analysis of parent genes of inherited circRNAs. (C) Location wise distribution of inherited circRNAs across chromosomes. The 25 chromosomes of zebrafish are scaled to the same length with zero representing the start of the chromosome and 1000 representing the end of the chromosome. The position of each inherited circRNA in the genome is represented by dots (Red).

To identify circRNA hotspots in the zebrafish genome, we checked expression of inherited circRNAs in zygote according to their position in the genome. The inherited circular RNAs are distributed across all chromosomes in zebrafish, with the maximum number (34) arising from chromosome 3, as seen in Figure 6A. Chromosomes 1, 5, 10 and 24 exhibit similar expression and distribution patterns across the length of chromosomes. A number of inherited circRNAs appear to be located near the ends of chromosomes, however, none of these were within the telomeric region (Figure 6C).

### Inherited circRNAs display stable expression till ZGA

The expression of circRNAs does not always correlate directly with the expression of their linear counterparts, as the back-splicing mechanism can be regulated separately from canonical splicing. We compared the expressions of inherited circRNAs and their linear counterparts in zygote. Overall, the inherited circRNA exhibit lower expression levels than their linear cognates (Figure 7A), consistent with the notion that circRNAs are expressed at low levels. Although a subset of these circRNAs is expressed at levels comparable to their linear counterparts, five of the inherited circRNAs show higher expression levels, with four of them originating from the same host gene (Supplementary file 10).

**Figure 7.**
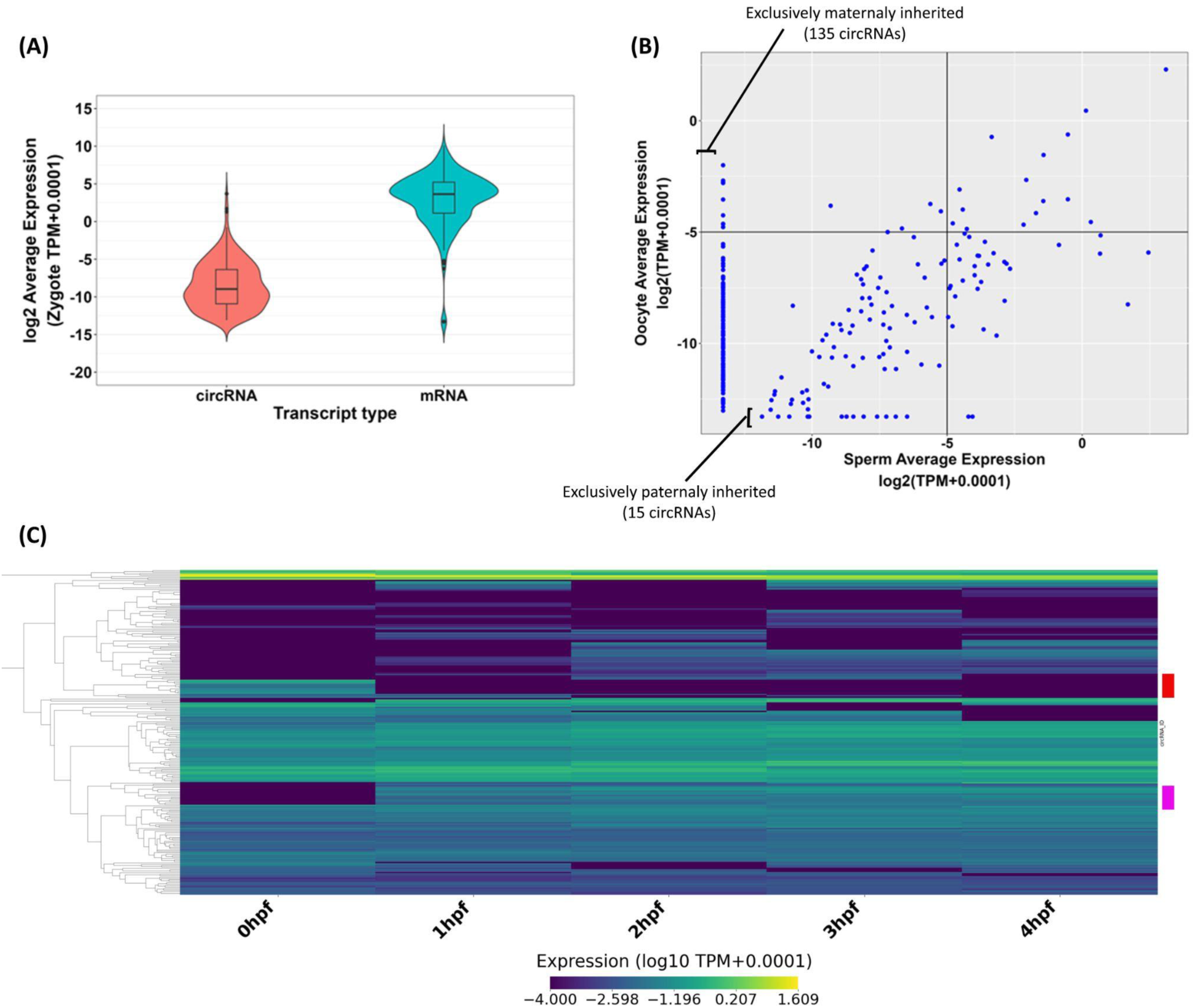
Inherited circRNA expression are mostly stable post fertilization till ZGA. (A) Violin plot showing the expression of all genic inherited circRNAs (231) and their corresponding mRNAs in the zygote. (B) Scatter plot showing the expression levels of 270 inherited circRNAs in the sperm and oocytes. C) Heatmap shows the expression dynamics of inherited circRNAs post fertilization. Each row is an individual circRNA expression. Two clusters are marked with red and pink blocks on the right, these show dynamic expression pattern during the early stages of development

Maternal mRNAs are known to be crucial for early development and are traditionally considered as the major contributor of RNAs prior to the zygotic genome activation (ZGA). To explore the parental origin of inherited circRNAs, we plotted the expression of the 270 inherited circRNA in sperm and oocytes. Unlike inherited lncRNAs, where >95% of them are present in both sperm and oocytes, only 45% percent of inherited circRNAs are present in both gametes and likely to be inherited from both parents. Half (135/270) of inherited circRNAs are exclusively expressed in oocytes. and only 5% are exclusively sperm derived. (Figure 7B; Supplementary file 11).

circRNAs are believed to be more stable and resistant to degradation due to their covalently closed loop structure. The absence of free 5’ and 3’ ends may protect them from exonucleases. To explore whether inherited circRNAs survive the global RNA degradation preceding zygotic genome activation, we plotted the temporal expression profiles of different clusters of circRNAs during the early developmental stages (0 hpf, 1 hpf, 2 hpf, 3 hpf and 4 hpf) (Figure 7C; Supplementary file 12) . We found that the majority of inherited circRNAs (∼70%) remained stable through the early developmental stages whether they were low or high expressors. We also found two interesting classes of inherited circRNA. The first group of inherited circRNAs were abundant at 0 hpf but expression dropped dramatically at 1 hpf and remained low till 4 hpf suggesting programmed degradation immediately after fertilization (Fig. 7C, red box). A second class of inherited circRNAs were expressed at very low levels at 0 hpf but the expression levels were elevated at 1 hpf and remained steady until 4 hpf (Fig. 7C, pink box). Thus, we observed stable expression in most inherited circRNAs but a small group showed quite dynamic expression patterns in early development (Figure 7C).

## Discussion

In this study we analyzed the inherited lncRNA and circRNA repertoire of zebrafish that consists of more than 2000 lncRNAs and more than 200 circRNAs representing ∼20% of all known lncRNAs and 7% of circRNAs in zebrafish. The actual number of inherited long regulatory RNAs could be even higher. In this study, the inherited lncRNA were identified by meta-analysis of published RNA sequencing datasets. This meta-analysis focused on consensus between multiple studies to shortlist only the high confidence inherited lncRNAs. The criteria that a lncRNA must be detected in multiple zygote and gametes RNA sequencing might have led to exclusion of some bonafide inherited lncRNAs. circRNAs, on the other hand, are rarer than lncRNAs, which themselves are not as abundant as mRNAs. Due to the lack of comprehensive annotation of circRNAs in zebrafish currently and the very low expression levels in the samples we used for analysis, we had to relax our criteria to include circRNAs that were expressed in at least one sample of sperm, oocyte or zygote. We have also used this less stringent criteria to annotate a circRNA as inherited. With improvements in genome annotation and RNA sequencing experiments at greater depth in future, we anticipate an even higher number of inherited long non-coding RNAs than currently shown in this study.

We find that the lncRNAs have a broad expression range in the zygote, with a minority of lncRNAs showing even higher expression than the median mRNA expression. Surprisingly, the sperm has similar diversity and expression of inherited lncRNAs as compared to the oocytes and lncRNA inheritance from sperm could be a novel theme in inheritance. However, an important point to ponder here is that the RNA sequencing libraries of sperm analyzed in this study are derived from thousands of spermatozoa and each individual spermatozoa may contain only a few copies of these lncRNAs. Whether this small number of copies of an lncRNA has a functional relevance or not remains to be seen.

We found more than 250 zebrafish inherited lncRNAs that have a conserved counterpart in the pool of both mouse and human inherited lncRNAs suggesting conserved functions for these inherited RNAs. We explored the functional relevance of inherited lncRNAs. Counterparts of these lncRNAs exist in many adult tissues at high levels. Thus, it does not seem that these lncRNAs are byproducts of oogenesis, inadvertently carried over during fertilization. An intellectually appealing, parsimonious hypothesis is that inherited RNAs may simply be a readily degradable source of nucleotides for the embryo. We looked at the post-fertilization expression pattern of inherited lncRNAs and circRNAs and found that most of these RNAs persist to 4 hpf, past the catastrophic RNA degradation and clearance of MZT. It appears that inherited lncRNAs and circRNAs are not merely a source of nucleotides or carry over of gametogenesis and they could have important RNA related functions in the zygote.

While inherited circRNAs exhibit a broad range of expression in the zygote, their expression levels are much lower than their linear counterparts, consistent with the general understanding that circRNAs are typically expressed at lower levels. A large subset of these circRNAs was predominantly expressed in oocytes, indicating their maternal origin. In the inherited circRNAs we found two classes of circRNAs that showed interesting expression patterns. A group of circRNAs appeared to be inherited in 0 hpf but were degraded rapidly by 1 hpf. This hints at a group of inherited circRNAs with functional relevance during fertilization. The second group was also interesting. From very low or no expression at 0 hpf these circRNAs appeared to be upregulated in 1hpf. Since it is known that no new transcription occurs in the zygote until the ZGA, around 4 hpf, this would suggest that there is active splicing happening in a small percentage of inherited linear RNA that might be giving rise to these circRNAs.

Modifications on the genomic DNA and histones are considered the major forms of epigenetic transfer of information from one generation to the next. Inheritance of non-coding regulatory RNAs, some of which have a long half-life, present an alternative pathway for transferring information from the parental generation to the offspring. Here, we provide a carefully annotated resource of inherited lncRNAs and circRNAs, along with information about their conservation and tissue specific expression. One of the surprising findings was the concordance between the inherited RNA pool in the male and female gametes. While maternal inheritance has been studied in a variety of organisms, the role of sperm derived RNA has been restricted to the recent reports of tRNA derived small RNA. In future, the functional studies on inherited lncRNAs and circRNAs can provide novel insights into various fields such as embryonic development, fertility and inheritance of acquired traits.

## Methods

### Meta-analysis of publicly available RNA sequencing datasets to identify inherited lncRNAs

We identified 2 samples (2 different studies) of sperm, 7 samples (4 different studies) of oocytes and 10 samples (4 different studies) of zygote RNA sequencing that were publicly available for zebrafish (Supplementary file 1). Similarly, we identified 4 sperm samples from 1 study and 4 oocytes and 2 zygote samples from another study for mouse (Supplementary file 1). For human, we identified 4 sperm samples from 1 study, 3 oocytes and 5 zygote samples from another study (Supplementary file 1). All these sequencing files were downloaded from the SRA (sequence read archive) of NCBI. The data was then analyzed using the bioinformatic pipeline outlined in Figure 1. Briefly, a quality check on all the RNA-seq datasets was performed using FastQC and low quality reads and adapters were trimmed using Trimmomatic-0.39 with following parameters - ILLUMINACLIP:adapters.fa:2:30:10 SLIDINGWINDOW:5:20 LEADING:3 TRAILING:3 MINLEN:50. The alignment of reads to the genome was performed using a splice-aware alignment tool STAR (version 2.7.1a) with the default parameters. The genome assemblies used for the zebrafish, mouse and human are GRCz11, GRCm38.p6 and GRCh38.p13 respectively. Expression quantification was done using the rsem-calculate-expression script of the RSEM package (version 1.3.1). For each species the resulting expression results (FPKM) were filtered for lncRNAs (i.e., the transcripts annotated as antisense, lncRNA, lincRNA, retained_intron, sense_intronic, sense_overlapping and processed_transcript in the Ensembl genome browser).

To identify inherited lncRNAs in zebrafish, firstly, the transcripts expressed (criteria - >=0.1 FPKM) in at least one sample of every zygote study were shortlisted. If the shortlisted transcripts from the above step were also expressed (criteria - >=0.1 FPKM) in at least one sample of any sperm or oocyte study then those transcripts were considered inherited. To identify the inherited lncRNAs in mouse and human, more stringent parameters were used as the samples are derived from only two publicly available studies. The lncRNAs expressed in the zygote (>=0.1 FPKM in all samples) and expressed in at least one of the gametes (>=0.1 FPKM in all samples) were considered inherited.

### Meta-analysis of publicly available RNA sequencing datasets to check the adult tissue expression profiles of inherited lncRNAs

For checking the expression of inherited lncRNAs in the adult tissues of zebrafish, we again used publicly available RNA sequencing datasets. We identified four studies that had profiled the expression in 15 adult tissues of zebrafish (Supplementary file 1). The tissues are brain, heart, muscle, liver, skin, gill, intestine, gut, testis, ovary, blood, spleen, bones, eye, kidney and tail. These samples were analyzed as per the analysis pipeline described above. Hierarchical clustering of the lncRNA expression profiles was performed using the linkage function from the scipy.cluster.hierarchy module.

### Conservation analysis

For sequence conservation analysis, the nucleotide sequences of 2093, 3785 and 4120 inherited lncRNAs in zebrafish, mouse and human respectively were extracted from the reference transcriptome using samtools faidx script. The zebrafish lncRNAs were searched for similarity in mouse and human sequences using stand-alone BLAST utility of NCBI. An E-value cutoff of 1e-5 and word size 20 was used to identify potentially sequence conserved lncRNAs, with no limit on query cover as regulatory elements can be embedded as small regions of similarity within a large lncRNA.

For identifying overlap conserved inherited lncRNAs, firstly, a list of conserved protein coding genes in zebrafish, mouse and human was obtained using biomart utility from Ensembl genome browser. Next, we extracted the genomic coordinates of inherited lncRNAs and conserved protein coding genes of zebrafish, mouse and human from their respective GTF files. Using Bedtools intersect script, inherited lncRNAs that have genomic overlap to conserved protein coding genes were shortlisted for each species. If a protein coding gene conserved in all three species also shows genomic overlap to inherited lncRNAs in all three species, then these lncRNAs were considered to be conserved by overlap.

For identifying loci conserved inherited lncRNAs, 10 neighboring protein coding genes for each inherited lncRNA were extracted (five each upstream and downstream) based on the genomic coordinates obtained from GTF files of zebrafish, mouse and human. Thus, corresponding to 2093, 3785 and 4120 inherited lncRNAs in zebrafish, mouse and human respectively, as many sets of 10 flanking protein coding genes were formed. These sets of flanking protein coding genes from zebrafish were compared with the sets from both mouse and human for conservation relationship. Zebrafish inherited lncRNAs with five or more flanking protein coding genes that have conserved counterparts in both mouse and human flanking protein coding gene sets were considered to be conserved by loci.

### GO analysis

The overlapping protein coding genes to genic inherited lncRNAs and the two flanking protein genes to intergenic lncRNAs were extracted from the GTF file. The parent genes to inherited circRNAs were identified by CIRI2 pipeline. To identify and categorize the biological processes enriched among these protein-coding genes ShinyGO 0.80 tool was used.

### Analysis of RNA sequencing data of post fertilization stage and clustering

The publicly available RNA sequencing data of 0 hpf, 1 hpf, 2 hpf, 3hpf and 4 hpf zebrafish embryos was reanalyzed using the pipeline described in the. The details of these samples are given in the table below (Supplementary file 1). The expression levels of 2093 inherited lncRNAs and 270 inherited circRNAs were extracted for the five stages. To classify the lncRNAs and circRNAs into distinct expression profiles, we applied the K-means clustering algorithm using the scikit-learn library in Python.

### Meta-analysis of publicly available RNA sequencing datasets to identify inherited circRNAs

We identified 2 samples (2 different studies) of sperm, 7 samples (4 different studies) of oocytes and 10 samples (4 different studies) of zygote RNA sequencing that were publicly available for zebrafish (Table 3-1). A brief schematic of the bioinformatic analysis is depicted in Figure 5. The raw RNA sequencing reads were subjected to a quality check using FastQC whereafter we discarded low quality reads and Illumina adapters using Trimmomatic (Bolger, Lohse, and Usadel 2014)(Bolger, Lohse et al. 2014, Gao, Zhang et al. 2017). For quantifying the expression of linear RNA, we aligned the processed RNA sequencing reads to the reference genome, zv11 using STAR aligner(Dobin et al. 2013) (Dobin, Davis et al. 2013). We used RNA-seq by Expectation-Maximization (RSEM)(B. Li and Dewey 2011) to quantify the expression of linear RNA transcripts and isoforms.

We used CIRI2 pipeline (Gao, Zhang, and Zhao 2018)(Gao, Zhang et al. 2017) with default settings to identify candidate circRNAs. The processed reads were aligned to the reference genome, zv11 using BWA aligner (H. Li and Durbin 2009) (Li and Durbin 2009). The output file from this alignment was used to annotate circRNAs using a Perl script available in CIRI2 package. We customised the perl script for our analysis, a circRNA having both coordinates outside any annotated gene boundaries was considered intergenic and it was considered as exonic or intronic if even one of the coordinates were located within annotated gene exon or intron. A candidate circRNA was called if it was supported by a minimum of 2 backsplice reads and at least a junction ratio of 0.05, which indicates towards the expression of a circRNA relative to its linear RNA. We then proceeded to identify inherited circRNA candidates. A circRNA was only considered to be inherited only if it was present in a zygotic sample along with either of the gametes. We identified 270 inherited circRNAs.

### Comparative analysis of circular RNAs and their linear isoforms

To investigate the host genes from which inherited circRNAs were transcribed, we annotated the circRNAs by linking their genomic positions to zebrafish gene loci. The analysis revealed that a single gene could produce multiple circRNA isoforms. Approximately six inherited circRNAs were found to span the coordinates of more than one gene. Among these, three circRNAs originated from overlapping genes with high sequence similarity. In contrast, two circRNAs aligned to distinct genes with dissimilar sequences. One circRNA, predicted to be approximately 145 kb in length, aligned with three distant genes. To compare circRNAs with their host RNAs, we analyzed the expression of each circRNA relative to the host genes to which they aligned. We quantified the expression of all linear RNAs in zebrafish using bioinformatics analysis (Figure 5A) and compared it with the normalized expression levels of each inherited circRNA.

## Acknowledgements

The work was supported by funding from Council for Scientific and Industrial Research (CSIR), India (project code: MLP2102). DCJ and AK were supported by research fellowships from CSIR and UGC respectively.

