## Supplementary material for "A comprehensive analysis of the inherited lncRNA and circRNA repertoire of zebrafish": Supplementary_files_caption.docx

Supplementary file 1- Details of publicly available RNA sequencing datasets used in this study.

Supplementary file 2- Details of 2093 inherited lncRNAs of zebrafish.

Supplementary file 3- List of inherited lncRNAs arising from ribosomal genes and hox loci.

Supplementary file 4- List of exclusively maternally and paternally inherited lncRNAs of zebrafish.

Supplementary file 5- List of inherited lncRNAs in 4 distinct groups based on their post-fertilization expression.

Supplementary file 6- Expression levels of inherited lncRNAs in zebrafish adult tissues.

Supplementary file 7- Details of mouse and human inherited lncRNAs.

Supplementary file 8- List of conserved inherited lncRNAs of zebrafish and their counterparts in mouse and human.

Supplementary file 9- Details of 270 inherited circRNAs of zebrafish.

Supplementary file 10- List of high expression inherited circRNAs of zebrafish.

Supplementary file 11- List of exclusively maternally and paternally inherited circRNAs of zebrafish.

Supplementary file 12- post-fertilization expression levels of inherited circRNAs.
